# Bacterial community composition and dynamics spanning five years in freshwater bog lakes

**DOI:** 10.1101/127035

**Authors:** Alexandra M. Linz, Benjamin C. Crary, Ashley Shade, Sarah Owens, Jack A. Gilbert, Rob Knight, Katherine D. McMahon

## Abstract

Bacteria play a key role in freshwater biogeochemical cycling, but long-term trends in freshwater bacterial community composition and dynamics are not yet well characterized. We used a multi-year time series of 16S rRNA gene amplicon sequencing data from eight bog lakes to census the freshwater bacterial community and observe annual and seasonal trends in abundance. Multiple sites and sampling events were necessary to begin to fully describe the bacterial communities. Each lake and layer contained a distinct bacterial community, with distinct levels of richness and indicator taxa that likely reflected the environmental conditions of each site. The community present in each year and site was also unique. Despite high interannual variability in community composition, we detected a core community of ubiquitous freshwater taxa. Although trends in abundance did not repeat annually, each freshwater lineage within the communities had a consistent lifestyle, defined by persistence, abundance, and variability. The results of our analysis emphasize the importance of long-term observations, as analyzing only a single year of data would not have allowed us to describe the dynamics and composition of these freshwater bacterial communities to the extent presented here.

**Importance:** Lakes are excellent systems for investigating bacterial community dynamics because they have clear boundaries and strong environmental gradients. The results of our research demonstrate that bacterial community dynamics operate on multi-year timescales, a finding which likely applies to other ecosystems, with implications for study design and interpretation. Understanding the drivers and controls of bacterial communities on long time scales would improve both our knowledge of fundamental properties of bacterial communities, and our ability to predict community states. In this specific ecosystem, bog lakes play a disproportionately large role in global carbon cycling, and the information presented here may ultimately help refine carbon budgets for these lakes. Finally, all data and code in this study are publicly available. We hope that this will serve as a resource to anyone seeking to answer their own microbial ecology questions using a multi-year time series.

## Introduction

One of the major goals of microbial ecology is to predict bacterial community composition. However, we have only a cursory knowledge of the factors that would allow us to predict bacterial community dynamics. To characterize the diversity and dynamics of an ecosystem’s bacterial community, sampling the same site multiple times is as necessary as sampling replicate ecosystems. Additionally, the sampling frequency must match the rate of change of the process being studied. We must first understand the scales on which bacterial communities change before we can design experiments that capture a full range of natural variation.

Bacterial communities have the potential to change more quickly than communities of macro-organisms due to their fast rate of reproduction. A meta-analysis of time series spanning one to three years found positive species-time relationships, indicating that more taxa are observed as the duration of sampling increases, either due to incomplete sampling, extinction and immigration, or speciation (1). Bacterial time series display time decay, meaning that the community continues to become more dissimilar from the initial sampling event as time from that event increases (2). In one freshwater lake, the amount of change in the bacterial community over a single day was equivalent to dissimilarity between sampling points ten meters apart (3). Conversely, bacterial communities can also change gradually over extremely long time scales, as they are sensitive to changes in environmental parameters such as nutrient availability and temperature. Wetland ecosystems and their carbon emissions are expected to change on scales greater than 300 years (4); as these emissions are the result of bacterial processes, we expect that the bacterial community will change on the same time scale as its ecosystem. Changes in marine phytoplankton regimes have been observed to occur over the past millennium, correlating with shifts in climate (5). With such a large range of potential change, we now recognize the need to more rigorously consider the duration and frequency of sampling in microbial ecology.

Multi-year studies of bacterial communities are less common due their logistical difficulties and the need for stable funding, but results from the United States National Science Foundation funded Microbial Observatory projects are exemplary. As a few examples among many, the San Pedro North Pacific - Microbial Observatory contributed to our understanding of heterogeneity of bacterial communities across space and time (6), while research at the Sapelo Island – Microbial Observatory has led the field in linking genomic data to metadata (7). In our own North Temperate Lakes – Microbial Observatory, based in Wisconsin, USA, a multi-year time series of metagenomic data was used to study sweeps in diversity at the genome level (8), adding to our knowledge of how genetic mutation influences bacterial communities. Long-term microbial ecology studies have a time-tested role in the quest to forecast bacterial communities.

Our North Temperate Lakes - Microbial Observatory time series was collected from eight bog lakes near Minocqua in the boreal region of northern Wisconsin, USA. Bog lakes contain high levels of dissolved organic carbon in the form of humic and fulvic acids, resulting in dark, “tea-colored” water. Due to their dark color, bog lakes absorb heat from sunlight, resulting in strong stratification during the summer. The top layer in a stratified bog lake, called the “epilimnion,” is oxygen-rich and warm. At the lake bottom, an anoxic, cold layer called the “hypolimnion” is formed. The transitions between mixing of these two layers and stratification occur rapidly in these systems, and at different frequencies (called mixing regimes) depending on the depth, surface area, and wind exposure of the lake. Changes in bacterial community composition along the vertical gradients established during stratification are well documented (9, 10). Mixing has been shown to be a disturbance to the bacterial communities in bog lakes (11). The bacterial communities in bog lakes are still being characterized, but contains both ubiquitous freshwater organisms (12, 13) and members of the candidate phyla radiation (14). Seasonality in these systems has been suggested (15, 16); however, multiple years of sampling are needed to confirm these prior findings.

Our dataset is comprised of 1,387 16S rRNA gene amplicon sequencing samples, collected from eight lakes and two thermal layers over five years. Our primary goal for this dataset was to census the bog lake community and determine which taxa are core to all bog lakes, to each thermal layer, and to each mixing regime. We also sought to learn how mixing regime structures the bacterial community, with our specific hypothesis being that lakes with intermediate levels of disturbance via mixing would be the most diverse. Finally, we investigated seasonality both at the community level and in individual taxa to identify annual trends. This extensive, long-term sampling effort establishes a time series that allows us to assess variability, responses to disturbance and re-occurring trends in freshwater bacterial communities.

## Results

### Overview of community composition

A time series of 16S amplicon data recovered from 1,387 samples was used to investigate bacterial community composition over time and across lakes. A total of 8,795 OTUs were detected. As is typical for most freshwater ecosystems, Proteobacteria, Actinobacteria, Bacteroidetes, and Verrucomicrobia were the most abundant phyla (Figure S1). Within these phyla, OTU abundance was highly uneven. For example, much of the abundance of *Proteobacteria* could be attributed to OTUs belonging to the well-known freshwater groups *Polynucleobacter* and *Limnohabitans*, and the freshwater clade acI contributed disproportionately to the observed abundance of Actinobacteria. Like many microbial communities, unevenness was a recurring theme in this dataset, which had a long rare tail of OTUs and trends driven largely by the most abundant OTUs (17, 18). Trimming of rare taxa did not impact the clustering observed in ordinations, such as those present in Figure 2, even when taxa observed less than 1000 times were removed.

### Community richness

We hypothesized that disturbance frequency, indicated by mixing regime, determines biodiversity levels. Observed richness was calculated for every sample at the OTU level, and samples were aggregated by lake and layer. Hypolimnia were generally richer than epilimnia (Figure 1, Table S1). Significant differences in richness between lakes were detected. For both layers, polymictic lakes had the fewest taxa, meromictic lakes had the most taxa, and dimictic lakes had intermediate numbers of taxa. This dataset includes two fall mixing events (Trout Bog 2007 and North Sparkling Bog 2008), as well as the artificial mixing event in North Sparkling Bog 2008 (11). Richness decreased sharply in mixed samples compared to those taken during the summer stratified period (Figure S2).

**Figure 1.**
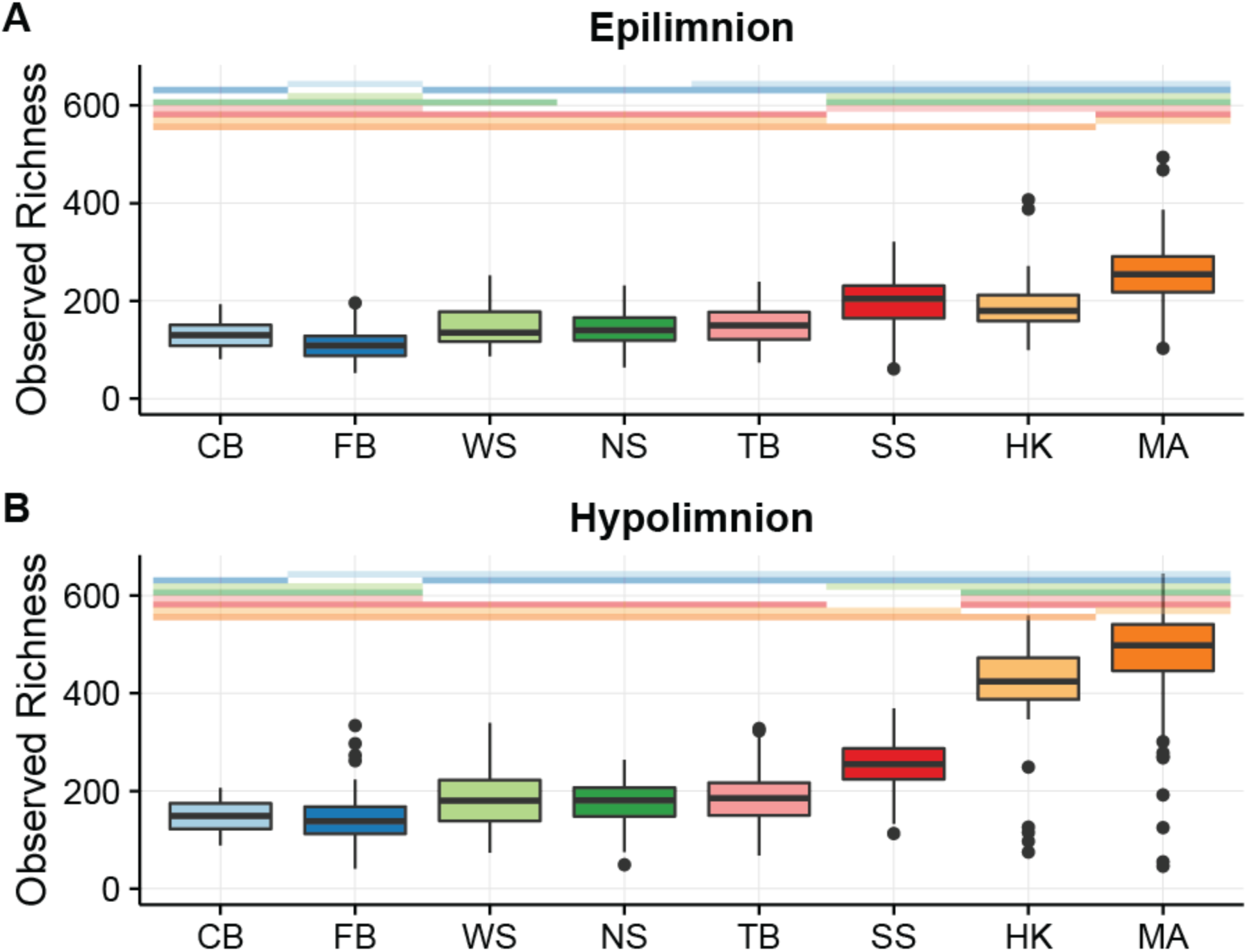
Richness by layer and lake. Lakes on the x axis are arranged by depth (see Table 1 for lake abbreviations and depth measurements). Lakes CB, FB, and WS are polymictic, lakes NS, TB, and SS are dimictic, and lakes HK and MA are meromictic. Colored bars above each plot represent significant differences in richness between lakes, with each colored bar matching the color of a lakes boxplot. For example, in Panel A, the boxplot for CB has the colored bars matching FB, NS, TB, SS, HK, and MA above it. This indicates that it is significantly different from these lakes, but not significantly different from the missing colored bar, WS. The mean and standard deviation for each lake and layer is reported in Table S1.

### Clusters of community composition

When differences in community composition were quantified using weighted UniFrac distance and visualized using principal coordinates analysis, several trends emerged. The large number of samples precluded much interpretation using a single PCoA, but sample clustering by layer, mixing regime, and lake was evident (Figure S3). Thus, we also examined PCoA for single lakes (both layers). Communities from the epilimnion and hypolimnion layers were significantly distinct from each other at p < 0.05 in all lakes except for polymictic Forestry Bog (FB) (p = 0.10) (Figure S4).

Within layers, mixing regime was the next factor explaining differences in community composition (Figure 2). Clustering by mixing regime was significant by PERMANOVA in both epilimnia and hypolimnia samples (r2 = 0.20 and r2 = 0.22, respectively, and p = 0.001 in both groups). Lake was a strong factor explaining community composition, with significant cluster in epilimnia (p = 0.001, r2 = 0.34) and hypolimnia (p = 0.001, r2 = 0.49).

**Figure 2.**
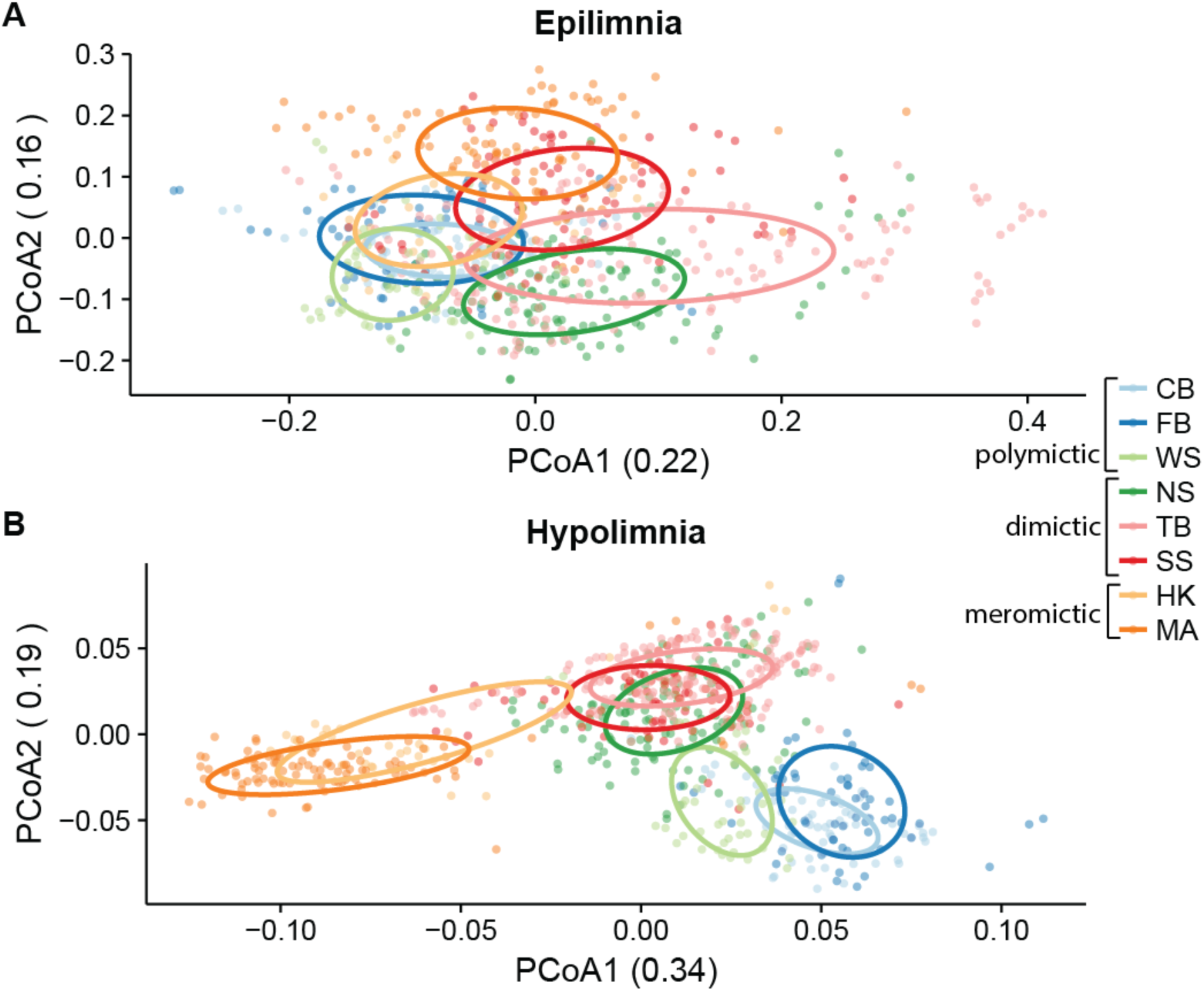
Weighted UniFrac distance was used to perform principal coordinates analysis on epilimnion (A) and hypolimnion (B) samples. In both layers, samples cluster significantly by lake and mixing regime as tested using PERMANOVA. (See Table 1 for lake abbreviations; CB, FB, and WS are polymictic, NS, TB, and SS are dimictic, HK and MA are meromictic). Ellipses indicating the clustering of each lake were calculate based on standard error using a 95% confidence interval. Differences in bacterial community composition between lakes and mixing regimes are more pronounced in hypolimnia than epilimnia.

### Variability and dispersion

While community composition was distinct by layer, lake, and mixing regime, there was still variability in community composition over time. Each year in each lake had a significantly different community composition, indicating interannual variability in the community composition (Figure 3a-c, Figure S5). We found no evidence of repeating seasonal trends during the stratified summer months in these lakes. Likewise, we examined the abundance trends of the most abundant individual OTUs and they did not seem to repeat each year, even when abundances in each year were normalized using z-scores (Figure S6).

**Figure 3.**
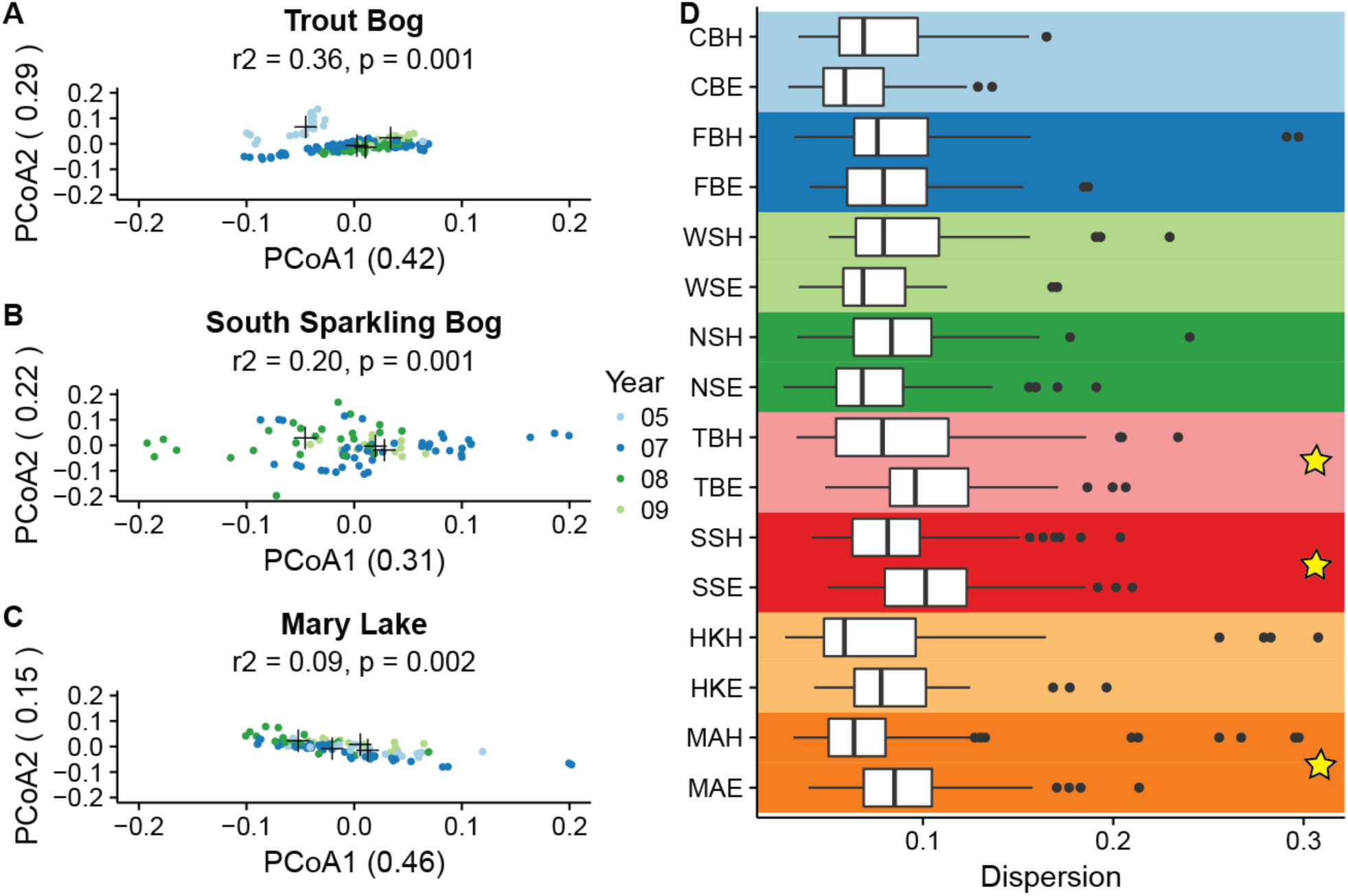
Internannual variability and dispersion by lake. Principal coordinates analysis using weighted UniFrac as the distance metric was used to measure the amount of interannual variation in the three lake hypolimnia with the longest time series (A-C). Additional ordinations of all other lakes and layers with at least two years of sampling are provided as supplemental figures (Figure S4). Black crosses indicated the centroid for each year. All hypolimnia showed significant clustering by year by PERMANOVA. Six outliers in Mary Lake from 2007 are not shown, as their coordinates lie outside the range specified for consistency between plots; these points were included in the PERMANOVA significance test. Panel D shows dispersion of each lake and layer in a PCoA including all samples (Lake abbreviations found in Table 1; E indicates epilimnion and H indicates hypolimnion; 6 outliers with distances from the centroid greater than 0.45 were removed). Stars indicate significant differences between layers at p < 0.05 by PERMADISP with a Bonferroni correction for multiple pairwise comparisons. Layers were signficantly different in TB, SS, and MA. No significant differences in dispersion between layers in the polymictic lakes (CB, FB, and WS), meromictic lake HK, or NS, a dimictic lake with an additional artificial mixing event.

Varibility can also be assessed by measuring the dispersion of groups in PCoA. Dispersion is the distance of each point from the centroid of a group on an ordination plot. This analysis showed that layers had significantly different degrees of dispersion in two of the dimictic lakes (Trout Bog and South Sparkling Bog) and a meromictic lake (Mary Lake) (Figure 3d). Two outliers in Mary Lake were removed; these dates showed different community compositions dominated by few taxa, possibily the result of a bloom event. Dispersion was not significantly different in the polymictic lakes, dimictic North Sparkling Bog, and meromictic Hell’s Kitchen. Increased sampling may reveal significant dispersion in these lakes. When dispersion between layers was significant, the epilimnion was on average more dispersed than the hypolimnion, indicating higher variability. This is consistent with our previously published results, and confirms that epilimnia are more variable than hypolimnia.

### The core community of bog lakes

One of the goals of this study was to determine the core bacterial community of bog lakes in general, and to determine if mixing regime affects core community membership. Our previous analyses showed that community composition was distinct in each layer and lake (Figure 2), while variability was observed within the same lake and layer (Figure 3). This prompted us to ask whether we had adequately sampled through time and space to fully census the lakes. Still, rarefaction curves generated for the entire dataset and for each layer begin to level off, suggesting that we have indeed sampled the majority of taxa found in our study sites (Figure S7). To identify the taxa that comprise the bog lake core community, we defined “core” as being present in 90% of a group of samples, regardless of abundance in the fully curated dataset. Four OTUs met this criteria for all samples in the dataset: OTU0076 (bacI-A1), OTU0097 (PnecC), OTU0813 (acI-B2), and OTU0678 (LD28). These taxa were therefore also core to both epilimnia and hypolimnia. Additional taxa core to epilimnia also included OTU0004 (betI), OTU0184 (acI-B3), OTU0472 (Lhab-A4), and OTU0522 (alfI-A1), while additional hypolimnia core taxa included OTU0042 (Rhodo), OTU0053 (unclassified Verrucomicrobia), and OTU0189 (acI-B2).

We performed the same core analysis after combining OTUs assigned to the same tribe (defined by 97% nucleotide similarity in the full length 16S region and phylogenetic branch structure (19)) into new groups. This revealed that certain tribes were core to the entire dataset or thermal layer even though their member OTUs were specific to certain sites. Notably, some OTUs were endemic to specific lakes, even though their corresponding tribe was found in multiple lakes/layers. OTUs not classified at the tribe level were not included. Results were similar to those observed at the OTU level, but yielded more core taxa. Tribes core to all samples included bacI-A1, PnecC, acI-B2, and LD28, but also betIII-A1 and acI-B4. In epilimnia, the core tribes were bacI-A1, PnecC, betIII-A1, acI-B3, acI-B2, Lhab-A4, alfI-A1, LD28, and acI-B4, while in hypolimnia, they were Rhodo, bacI-A1, PnecC, betIII-A1, acI-B2, and acI-B4. These results show that despite lake-to-lake differences and interannual variability, there are bacterial taxa that are consistently present in bog lakes.

Principal coordinates analysis suggested that samples clustered also by mixing regime (Figure 2). We thus evaluated Venn diagrams of OTUs shared by and unique to each mixing regime to better visualize the overlap in community composition (Figure 4). In both epilimnia and hypolimnia, meromictic lakes had the greatest numbers of unique OTUs while polymictic lakes had the least, consistent with the differences in richness between lakes (Figure 1). Shared community membership, i.e. the number of OTUs present at any abundance in both communities, differed between mixing regimes. Epilimnia (A) and hypolimnia (B) showed similar trends in shared membership: meromictic and dimictic lakes shared the most OTUs, while meromictic and polymictic lakes shared the least.

**Figure 4.**
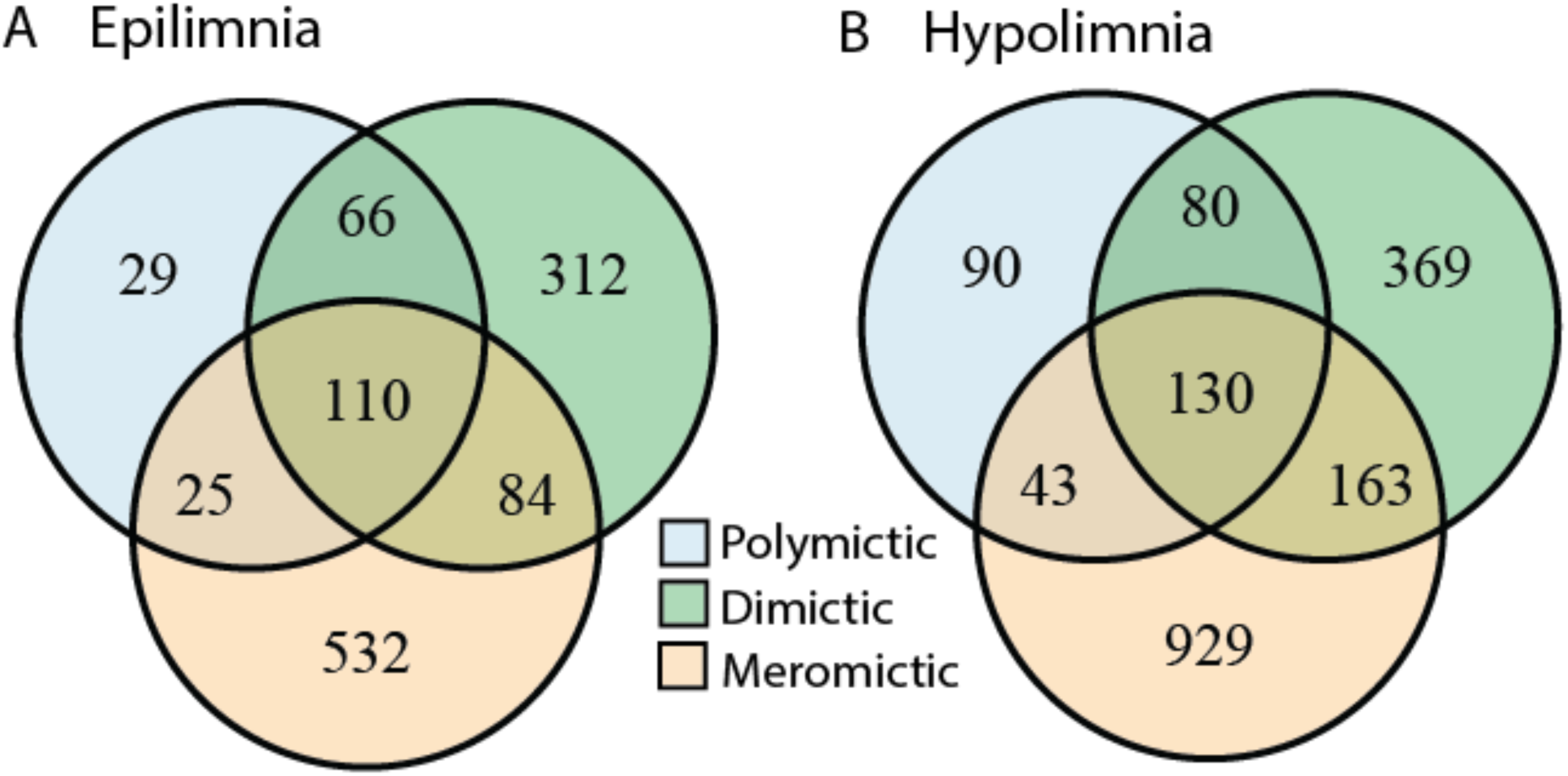
Numbers of unique and shared OTUs by mixing regime. To better understand how shared community membership differs by mixing regime, we quantified the number of shared and unique OTUs in each category. An OTU needed only to appear in one sample at any abundance to be considered present in a category. We found that in both layers, meromictic lakes have the greatest numbers of unique OTUs and polymictic lakes have the least. Meromictic and dimictic lakes shared the most OTUs, while meromictic and polymictic lakes shared the least. Dimictic lakes shared more OTUs with meromictic lakes than with polymictic lakes.

We next used indicator analysis to identify the taxa unique to each mixing regime. Indicator analysis is a statistical method used to determine if taxa are found significantly more frequently in certain pre-determined groups of samples than in others. In this case, the groups were defined by mixing regime, and normalization was applied to account for different numbers of samples in each group. OTUs were grouped at every taxonomic level, and all taxonomic levels were run in the indictor analysis at once to account for differences in the ability of these levels to serve as indicators (for example, the order Actinomycetales is a stronger indicator of polymictic lakes than the phylum Actinobacteria). An abundance threshold of 500 reads was imposed on each taxonomic group. The full table of results from the indicator analysis is available in the supplemental material, while a few indicator taxa of interest are highlighted here.

The lineage acI is a ubiquitous freshwater group, with specific clades and tribes showing a preference for bog lakes in previous studies (20, 21). Our dataset shows a further distinction of acI by mixing regime in epilimnia; acI-A tribes were found predominantly in meromictic lakes, with exception of Phila, which is an indicator of polymictic lakes. Tribes of acI-B, particularly OTUs belonging to acI-B2, were indicators of dimictic lakes. Methylophilales, a putative methylotroph, was also an indicator of dimictic lakes, as was putative sulfate reducer Desulfobulbaceae. The phyla Planctomyces, Omnitrophica (formerly OP3), OP8, and Verrucomicrobia were found more often in meromictic lakes, as were putative sulfate reducers Syntrophobacterales and Desulfobacteraceae. Indicators of polymictic lakes include ubiquitous freshwater groups such as Limnohabitans, Polynucleobacter (PnecC), betI-A, and verI-A. These indicator taxa likely reflect the environmental conditions unique to each mixing regime.

### Lifestyles of freshwater lineages

Even though OTUs do not show the same trends each year, they do possess patterns that are consistent between years and lakes. We quantified mean abundance when present, persistence (defined as the proportion of samples containing the group of interest), and the coefficient of variation for lineages classified using the freshwater taxonomy, metrics which have been previously used to categorize OTUs (22, 23). Using only these well-defined freshwater groups allowed better taxonomic resolution as we summed the abundances of OTUs by their lineage classification. Lifestyle traits of lineages were consistent across both lakes and years. Low persistence was associated with high variability, and low variability was associated with high abundance (Figure 5, Figure S8). We rarely observed “bloomers,” situations where a clade had both high abundance and low persistence; one potential reason for this could be that true “bloomers” drop below the detection limit of our sequencing methods when not abundant. Most freshwater lineages were highly persistent at low abundances with low variability. Lineage gamIII of the Gammaproteobacteria was an exception, with low persistence, low abundance, and high variability. Lineages gamI and verI-A occasionally also exhibited this profile. Lineages betII and acI were highly abundant and persistent with low variability, consistent with their suggested lifestyles as ubiquitous freshwater generalists (12, 21).

**Figure 5.**
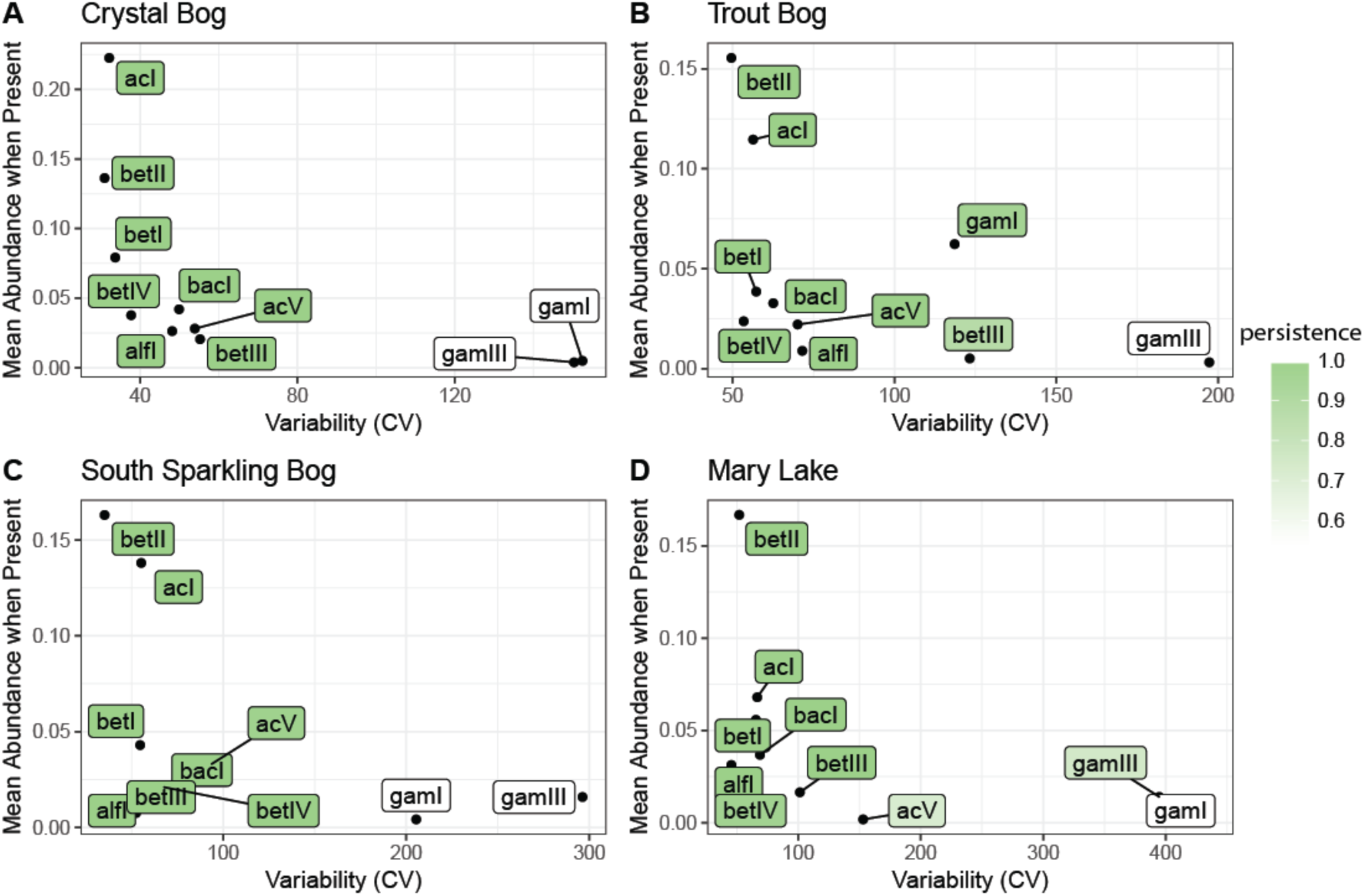
Traits of freshwater lineages. These well-defined freshwater groups showed similar persistence, variance, and abundance in every lake, despite differing abundance patterns. Data from epilimnia with at least two years of undisturbed sampling are shown here. Mean abundance was represented as the average percentage of reads attributed to each lineage when that lineage was present. Variability was measured as the coefficient of variation. Persistence (shaded color) was defined as the proportion of samples containing each lineage. The observed consistency in mean abundance, variability, and persistence suggests that unknown functions or metabolic characteristics drive a stable lifestyle. Additional plots by year can be found in Figure S8.

## Discussion

The North Temperate Lakes - Microbial Observatory dataset is a comprehensive 16S amplicon survey spanning four years, eight lakes, and two thermal layers. We found that multiple years of sampling were necessary to census the community of bog lake ecosystems. Richness and membership in these communities were structured by layer, mixing regime, and lake. We identified specific bacterial taxa present throughout the dataset, as well as taxa endemic to certain depths or mixing regimes. Mixing events were associated with reduced richness and an increase in the proportion of certain taxa. High levels of variability were detected in this dataset; each year in each lake harbored a unique bacterial community. However, freshwater lineages still showed consistent lifestyles, defined by abundance, persistence, and variability, across lakes and years, even though the abundance trends of individual populations were. Our results emphasize the importance of multiple sampling events to assess full bacterial community membership and variability in an ecosystem.

The bog lakes in this study have been model systems for freshwater microbial ecology for many years. Early studies used Automated Ribosomal Intergenic Spacer Analysis (ARISA), a fingerprinting technique for identifying unique bacterial taxa in environmental samples (24). Our research built upon these studies and added information about the taxonomic identities of bacterial groups. For example, persistent and unique bacterial groups were detected in the bog lakes using ARISA (25); using 16S amplicon sequencing, we also found persistent groups and could identify them as the ubiquitous freshwater bacteria LD28, acI-B2, PnecC, and bacI-A1. Differences in richness and community membership were previously detected between Crystal Bog, Trout Bog, and Mary Lake, three sites representative of the three mixing regime categories of polymictic, dimictic, and meromictic (25). Our data supported these results and suggest that these trends are indeed linked with mixing regime, as we included multiple lakes of each type sampled over multiple years in this study.

We also supported previous research on the characteristics of bacterial communities in the epilimnion and hypolimnion, and the impacts of lake mixing on these communities. We confirmed that epilimnia communities tended to be more dispersed than hypolimnia communities, potentially due to increased exposure to climatic events (25). Mixing was disruptive to both epilimnion and hypolimnion communities, selecting for only a few taxa that persist during this disturbance, but quickly recovering diversity once stratification was re-established (11, 26). Comparing richness between lakes of different mixing regimes did not support the intermediate disturbance hypothesis, which was our initial inspiration for the collection of this dataset; rather, the least frequently mixing lakes had the most diverse communities. As many variables co-vary with mixing regime (such as depth, volume of integrated water column, dissolved carbon concentrations and total nitrogen concentration), it is not clear which variables are driving this trend. One likely explanation is that increased depth leads to increased habitat heterogeneity, as more distinct niches develop along the vertical chemical gradients in the lake. These results are in line with current thinking in the ecological community, as other studies on diversity and disturbance have also found no evidence for the intermediate disturbance hypothesis (27).

We were not able to detect repeatable annual trends in bog lakes in our multiple years of sampling. While seasonality in marine and river systems has been well-established by our colleagues, previous research on seasonality in freshwater lakes has produced inconsistent results (28–31). Distinct, seasonally repeatable community types were identified in alpine lakes, but stratified summer communities were distinct each year (32). Seasonal trends were detected in a time series from Lake Mendota similar to this study, but summer samples in Lake Mendota were more variable then those collected in other seasons (33). In the previous ARISA-based research on the bog lakes in our dataset, community properties such as richness and rate of change were consistent each year, and the phytoplankton communities were hypothesized to drive seasonal trends in the bacterial communities based on correlation studies (34–36). Synchrony in seasonal trends was observed (35); however, in a second year of sampling for seasonal trends in Crystal Bog and Trout Bog, these findings were not reproduced (37). Successional trends were studied in Crystal Bog and Lake Mendota with a relatively small number of samples collected over two years and “dramatic changes” in community composition associated with drops in biodiversity were described during the summer months, while spring, winter, and fall had more stable community composition (34). Because our dataset was sparsely represented by seasons other than summer, higher summer variability may explain why we see a different community each year and a lack of seasonal trends in community composition. However, we cannot disprove the influence of seasonality on bacterial community dynamics in temperate freshwater lakes.

One of the biggest benefits of 16S rRNA gene amplicon sequencing over ARISA is the ability to assign names to sequences. In addition to a core of persistent taxa found in nearly every sample collected, we also identified taxa endemic to either the epilimnion or hypolimnion and to specific mixing regimes. These endemic taxa likely reflect the biogeochemical differences driven by mixing regime. Dimictic and meromictic hypolimnia, which are consistently anaerobic, harbor putative sulfur and sulfate reducers not present in polymictic hypolimnia, which are more frequently oxygenated. Members of the acI lineage partition by mixing regime in epilimnia, and the functional traits driving this filtering effect are the subject of active study (20). Interestingly, the meromictic Mary Lake hypolimnion contains several taxa classified into the candidate phyla radiation and a larger proportion of completely unclassified reads than other hypolimnia (38). This is consistent with the findings of other 16S and metagenomics studies of meromictic lakes (39, 40), and suggests that the highly reduced and consistently anaerobic conditions in meromictic hypolimnia are excellent study systems for research on members of the candidate phyla radiation and “microbial dark matter”.

Perhaps the biggest implication of this study is the importance of repeated sampling of microbial ecosystems. A similar dataset spanning only a single year would not have captured the full extent of variability observed, and therefore would not have detected as many of the taxa belonging to the bog lake community; even our four years of weekly sampling did not result in level rarefaction curves (Figure S7). While we found no evidence for seasonal trends or repeated annual trends, it is possible that there are cycles or variables acting on scales longer than the five years covered in this dataset, or that annual differences are driven by environmental factors that do not occur every year. Unmeasured biotic interactions between bacterial taxa may also contribute to the observed variability. Understanding the factors that contribute to variability in bog lake communities will lead to improved predictive modelling in freshwater systems, allowing forecasting of bloom events and guiding better management strategies. Additionally, these systems may be ideal for addressing some of the core questions in microbial ecology, such as how community assembly occurs, how interactions between taxa shape community composition, and how resource partitioning drives the lifestyles of bacterial taxa.

To answer these questions and more, we continue to collect and sequence samples for the North Temperate Lakes – Microbial Observatory, and we are expanding our sequencing repertoire beyond 16S rRNA gene sequencing. All 16S rRNA gene data we have currently generated can be found in the R package “OTUtable” which is available on CRAN for installation via the R command line, or on our GitHub page. This dataset has already been used in a meta-analysis of microbial time series (1). We hope that this dataset and its future expansion will be used as a resource for researchers investigating their own questions about how bacterial communities behave on long time scales.

## Materials and Methods

### Sample Collection

Water was collected from eight bog lakes during the summers of 2005, 2007, 2008 and 2009, as previously described (25). Briefly, the epilimnion and hypolimnion layers were collected separately using an integrated water column sampler. Dissolved oxygen and temperature profiles were measured at the time of collection using a handheld YSI 550A (YSI Inc., Yellow Springs, OH). After transport to the laboratory, approximately 150 mL from each well-mixed sample was filtered through a 0.22 micron polyethersulfone filter (Supor 200, Pall, Port Washington, NY). Filters were stored at -80C until DNA extraction using FastDNA Spin Kit for Soil (MP Biomedicals, Santa Ana, CA), with minor modifications (41). The sampling sites are located near Boulder Junction, WI, and were chosen to include lakes represent the three mixing regimes of polymictic (multiple mixing events per year), dimictic (two mixing events per year, usually in spring and fall), and meromictic (no record mixing events) (Table 1). Trout Bog and Crystal Bog are also primary study sites for the North Temperate Lakes - Long Term Ecological Research Program, which measures a suite of chemical limnology parameters fortnightly during the open water season. The NTL-LTER also maintains autonomous sensing buoys on Trout Bog and Crystal Bog, allowing for more refined mixing event detection based on thermistor chain measurements.

**Table 1.** Location and characteristics of study sites. The lakes included in this time series are small, humic bog lakes in the boreal region near Minocqua, Wisconsin, USA. They range in depth from 2 to 21.5 meters and encompass a range of water column mixing frequencies (termed regimes). Dimictic lakes mix twice per year, typically in fall and spring, while polymictic lakes can mix more than twice throughout the spring, summer, and fall. Meromictic lakes have no recorded mixing events. pH was measured in 2007, while nutrient data was measured in 2008 (with the exceptions of FB, WS, and HK, measured in 2007). When two values are present in a single box, the first represents the epilimnion value and the second represents the hypolimnion value.

|  | **Forestry Bog** | **Crystal Bog** | **North Sparkling Bog** | **West Sparkling Bog** | **Trout Bog** | **South Sparkling Bog** | **Hell's Kitchen** | **Mary Lake** |
| --- | --- | --- | --- | --- | --- | --- | --- | --- |
| **ID** | FB | CB | NS | WS | TB | SS | HK | MA |
| **Depth (*m*)** | 2.0 | 2.5 | 4.5 | 4.6 | 7.0 | 8.0 | 19.3 | 21.5 |
| **Surface area (*m*<sup>2</sup>)** | 1300 | 5600 | 4700 | 11900 | 10100 | 4400 | 30000 | 12000 |
| **Mixing regime** | Polymictic | Polymictic | Dimictic | Polymictic | Dimictic | Dimictic | Meromictic | Meromictic |
| **GPS coordinates** | 46.04777, -89.651248 | 46.007639, -89.606341 | 46.004819, -89.705214 | 46.004633, -89.709082 | 46.041140, -89.686352 | 46.041140, -89.709082 | 46.186674, -89.702510 | 46.250764, -89.900419 |
| **Years sampled** | 2007 | 2007, 2009 | 2007, 2008, 2009 | 2007 | 2005, 2007, 2008, 2009 | 2007, 2008, 2009 | 2007 | 2005, 2007, 2008, 2009 |
| **pH** | 4.97, 4.85 | 4.49, 4.41 | 4.69, 4.80 | 5.22, 5.14 | 4.60, 4.78 | 4.46, 4.94 |  | 5.81, 5.72 |
| **Dissolved inorganic carbon (ppm)** | 0.94, 1.46 | 0.69, 1.72 | 1.12, 2.31 | 0.76, 1.56 | 1.73, 4.47 | 1.97, 6.42 | 2.91, 9.70 | 5.54, 12.38 |
| **Dissolved organic carbon (ppm)** | 10.22, 8.96 | 15.47, 13.6 | 10.05, 10.40 | 7.26, 7.27 | 19.87, 20.58 | 12.40, 21.92 | 7.26, 7.33 | 20.63, 67.10 |
| **Total nitrogen (ppb)** |  | 620.57, 846.00 | 629.09, 809.45 |  | 737.71, 1121.00 | 813.88, 1498 |  | 1332.57, 3652.38 |
| **Total phosphorus (ppb)** |  | 30.00, 38.86 | 78.00, 135.45 |  | 50.57, 53.25 | 48.63, 69.14 |  | 78.00, 303.50 |
| **Total dissolved nitrogen (ppb)** |  | 1290.19, 490.13 | 442.39, 586.56 |  | 582.5, 820.21 | 451.63, 1179.21 |  | 1024.5, 3220.14 |
| **Total dissolved phosphorus (ppb)** |  | 84.25, 14.88 | 70.22, 22.67 |  | 34.5, 31.57 | 16.25, 18.29 |  | 71.13, 228 |

### Sequencing

A total of 1,510 DNA samples, including 547 biological replicates, were sequenced by the Earth Microbiome Project according to their standard protocols in 2010, using the original V4 primers (FWD:GTGCCAGCMGCCGCGGTAA;REV:GGACTACHVGGGTWTCTAAT) (42). Briefly, the V4 region was amplified and sequenced using Illumina HiSeq, resulting in 77,517,398 total sequences with an average length of 150 base pairs. To reduce the number of erroneous sequences, QIIME’s “deblurring” algorithm for reducing sequence error in Illumina data was applied (43). Based on the sequencing error profile, this algorithm removes reads that are likely to be sequencing errors if those reads are both low in abundance and highly similar to a high abundance read. Reads occurring less than 25 times in the entire dataset were removed after deblurring, leaving 9,856 unique sequences. These sequences are considered operational taxonomic units (OTUs).

570 sequences with long homopolymer runs, ambiguous base calls, or incorrect sequence lengths were found and removed via mothur v1.34.3 (44). Thirty-three chimeras and 340 chloroplast sequences (based on pre-clustering and classification with the Greengenes 16S database, May 2013) (45) were removed. Samples were rarefied to 2,500 reads; samples with less than 2,500 reads were omitted, resulting in 1,387 remaining samples. The rarefaction cutoff used was determined based on the results of simulation; 2,500 reads was chosen to maximize the number of samples retained, while maintaining sufficient quality for downstream analysis of diversity metrics.

Representative sequences for each OTU were classified in either our curated freshwater database (19) or the Greengenes database based on the output of NCBI-BLAST (blast+ 2.2.3.1) (46). Representative sequences from each OTU were randomly chosen. The program blastn was used to compare representative sequences to full-length sequences in the freshwater database. OTUs matching the freshwater database with a percent identity greater than 98% were classified in that database, and remaining sequences were classified in the Greengenes database. Both classification steps were performed in mothur using the Wang method (47), and classifications with less than 70% confidence were not included. A detailed workflow for quality control and classification of our sequences is available at (https://github.com/McMahonLab/16STaxAss ) (manuscript in prep).

### Statistics

Statistical analysis was performed in R v3.3.2 (R Development Core Team, 2008. R: A language and environment for statistical computing.). Significant differences in richness between lakes was tested using a pairwise Wilcoxon sum rank test with a Bonferroni adjustment in the R package “exactRankTests” (T. Hothorn and K. Hornik, 2015. exactRankTests: Exact Distributions for Rank and Permutation Tests). Similarity between samples was compared using UniFrac distances, as implement in “phyloseq” (48) (P.J. McMurdie and S. Holmes, 2013. phyloseq: An R Package for reproducible interactive analysis and graphic of microbiome census data). Weighted and unweighted Unifrac distance (48) was compared with Bray-Curtis Dissimilarity and Jaccard Similarity, implemented in “vegan” (J. Oksanen, 2016. vegan: Community Ecology Package). Weighted UniFrac distances were chosen for principle coordinates analysis, performed by betadisper() in “vegan”, because it explained the greatest amount of variation in the first two axes. Significant clustering by year in PCoA and in dispersion between lakes was tested using PERMADISP with the function adonis() in “vegan.”

Indicator species analysis was performed using “indicspecies” (49). Only taxa with read abundances of at least 500 reads in the entire dataset were used for this analysis. The group-normalized coefficient of correlation was chosen for this analysis because it measures both positive and negative habitat preferences and accounts for differences in the number of samples from each site. All taxonomic levels were included in this analysis to determine which level of resolution was the best indicator for each taxonomic group.

Plots were generated using “ggplot2” (H. Wickham, 2009. ggplot2: Elegant Graphics for Data Analysis) and “cowplot” (C. Wilke, 2016. cowplot: Streamlined Plot Themes and Plot Annotations for ‘ggplot2’). “reshape2” was used for data formatting (H. Wickham, 2007. Reshaping Data with the reshape Package). Data and code from this study can be downloaded from the R package “OTUtable” and the McMahon Lab GitHub repository “North_Temperate_Lakes-Microbial_Observatory.”

## Acknowledgments

This study was made possible through generous support by the Earth Microbiome Project, which is funded in part through awards by the Keck and Templeton Foundations. AL is supported by the National Science Foundation Graduate Research Fellowship Program under Grant No. (DGE-1256259). KDM acknowledges funding from the United States National Science Foundation Microbial Observatories program (MCB-0702395), the Long Term Ecological Research program (NTL-LTER DEB-1440297) and an INSPIRE award (DEB-1344254).

We thank the North Temperate Lakes Microbial Observatory 2005, 2007, 2008, and 2009 field crews, UW-Trout Lake Station, the UW Center for Limnology, and the Global Lakes Ecological Observatory Network for field and logistical support. Special thanks to Sara Paver and Sara Yeo for coordinating field crews in 2009. We thank Greg Caporaso for contributions during early stages of data analysis and Amnon Amir for early access to the deblurring algorithm. We thank McMahon Lab members Robin Rohwer and Joshua Hamilton for early access to a workflow used to assign taxonomies to OTUs using a custom 16S training set. We acknowledge efforts by many McMahon Lab undergrads and technicians related to sample collection and DNA extraction, particularly Georgia Wolfe.

## References

1. Shade A, Caporaso JG, Handelsman J, Knight R, Fierer N. 2013. A meta-analysis of changes in bacterial and archaeal communities with time. ISME J 7: 1493–506.

2. Faust K, Lahti L, Gonze D, de Vos WM, Raes J. 2015. Metagenomics meets time series analysis: unraveling microbial community dynamics. Curr Opin Microbiol 25: 56–66.

3. Jones SE, Cadkin TA, Newton RJ, McMahon KD. 2012. Spatial and temporal scales of aquatic bacterial beta diversity. Front Microbiol 3: 1–10.

4. Mitsch WJ, Bernal B, Nahlik AM, Mander Ü, Zhang L, Anderson CJ, Jørgensen SE, Brix H. 2013. Wetlands, carbon, and climate change. Landsc Ecol 28: 583–597.

5. McMahon KW, McCarthy MD, Sherwood OA, Larsen T, Guilderson TP. 2015. Millennial-scale plankton regime shifts in the subtropical North Pacific Ocean. Science (80- ) 350: 1530–1533.

6. Hewson I, Steele JA, Capone DG, Fuhrman JA. 2006. Remarkable heterogeneity in meso- and bathypelagic bacterioplankton assemblage composition. Limnol Oceanogr 51: 1274–1283.

7. Gifford SM, Sharma S, Moran MA. 2014. Linking activity and function to ecosystem dynamics in a coastal bacterioplankton community. Front Microbiol 5: 1–12.

8. Bendall ML, Stevens SLR, Chan L, Malfatti S, Schwientek P, Tremblay J, Schackwitz W, Martin J, Pati A, Bushnell B, Froula J, Kang D, Tringe SG, Bertilsson S, Moran MA, Shade A, Newton RJ, McMahon KD, Malmstrom RR. 2016. Genome-wide selective sweeps and gene-specific sweeps in natural bacterial populations. ISME J 10: 1589–1601.

9. Taipale S, Jones R, Tiirola M. 2009. Vertical diversity of bacteria in an oxygen-stratified humic lake, evaluated using DNA and phospholipid analyses. Aquat Microb Ecol 55: 1–16.

10. Garcia SL, Salka I, Grossart HP, Warnecke F. 2013. Depth-discrete profiles of bacterial communities reveal pronounced spatio-temporal dynamics related to lake stratification. Environ Microbiol Rep 5: 549–555.

11. Shade A, Read JS, Youngblut ND, Fierer N, Knight R, Kratz TK, Lottig NR, Roden EE, Stanley EH, Stombaugh J, Whitaker RJ, Wu CH, McMahon KD. 2012. Lake microbial communities are resilient after a whole-ecosystem disturbance. ISME J 6: 2153–2167.

12. Hahn MW, Scheuerl T, Jezberová J, Koll U, Jezbera J, Šimek K, Vannini C, Petroni G, Wu QL. 2012. The passive yet successful way of planktonic life: Genomic and experimental analysis of the ecology of a free-living polynucleobacter population. PLoS One 7: 1–17.

13. Garcia SL, McMahon KD, Grossart HP, Warnecke F. 2013. Successful enrichment of the ubiquitous freshwater acI Actinobacteria. Environ Microbiol Rep 1–7.

14. Peura S, Eiler A, Bertilsson S, Nyka H, Tiirola M, Jones RI. 2012. Distinct and diverse anaerobic bacterial communities in boreal lakes dominated by candidate division OD1. ISME J J 6: 1640–1652.

15. Eiler A, Heinrich F, Bertilsson S. 2012. Coherent dynamics and association networks among lake bacterioplankton taxa. ISME J 6: 330–42.

16. Graham JM, Kent AD, Lauster GH, Yannarell AC, Graham LE, Triplett EW. 2004. Seasonal dynamics of phytoplankton and planktonic protozoan communities in a northern temperate humic lake: Diversity in a dinoflagellate dominated system. Microb Ecol 48: 528–540.

17. Mariadassou M, Pichon S, Ebert D. 2015. Microbial ecosystems are dominated by specialist taxa. Ecol Lett 18: 974–982.

18. Newton RJ, Shade A. 2016. Lifestyles of rarity : understanding heterotrophic strategies to inform the ecology of the microbial rare biosphere. Aquat Microb Ecol 78: 51–63.

19. Newton RJ, Jones SE, Eiler A, McMahon KD, Bertilsson S. 2011. A guide to the natural history of freshwater lake bacteria. Microbiol Mol Biol Rev 75: 14–49.

20. Garcia SL, Buck M, McMahon KD, Grossart H-P, Eiler A, Warnecke F. 2015. Auxotrophy and intra-population complementary in the “interactome” of a cultivated freshwater model community. Mol Ecol 24: 4449–4459.

21. Ghylin TW, Garcia SL, Moya F, Oyserman BO, Schwientek KT, Mutschler J, Dwulit-Smith J, Chan L-K, Martinez-Garcia M, Sczyrba A, Stepanauskas R, Grossart H-P, Woyke T, Warnecke F, Malmstrom R, Bertilsson S, McMahon KD. 2014. Comparative single-cell genomics reveals potential ecological niches for the freshwater acI Actinobacteria lineage. ISME J 8: 2503–2516.

22. Herren CM, Webert KC, McMahon KD. 2016. Environmental Disturbances Decrease the Variability of Microbial Populations within Periphyton. mSystems 1: 1–14.

23. Shade AL, Gilbert JA. 2015. Temporal patterns of rarity provide a more complete view of microbial diversity. Trends Microbiol 23: 335–340.

24. Fisher MM, Triplett EW. 1999. Automated approach for ribosomal intergenic spacer analysis of microbial diversity and its application to freshwater bacterial communities. Appl Environ Microbiol 65: 4630–4636.

25. Shade A, Jones SE, McMahon KD. 2008. The influence of habitat heterogeneity on freshwater bacterial community composition and dynamics. Environ Microbiol 10: 1057–1067.

26. Shade A, Read JS, Welkie DG, Kratz TK, Wu CH, McMahon KD. 2011. Resistance, resilience and recovery: Aquatic bacterial dynamics after water column disturbance. Environ Microbiol 13: 2752–2767.

27. Fox JW. 2013. The intermediate disturbance hypothesis should be abandoned. Trends Ecol Evol 28: 86–92.

28. Crump BC, Hobbie JE. 2005. Synchrony and seasonality in bacterioplankton communities of two temperate rivers. Limnol Oceanogr 50: 1718–1729.

29. Gilbert JA, Steele JA, Caporaso JG, Steinbruck L, Reeder J, Temperton B, Huse S, McHardy AC, Knight R, Joint I, Somerfield P, Fuhrman JA, Field D. 2012. Defining seasonal marine microbial community dynamics. ISME J 6: 298–308.

30. Fuhrman JA, Hewson I, Schwalbach MS, Steele JA, Brown M V, Naeem S. 2006. Annually reoccurring bacterial communities are predictable from ocean conditions. Proc Natl Acad Sci USA 103: 13104–13109.

31. Cram J, Chow C, Sachdeva R, Needham D, Parada A, Steele J, Fuhrman J. 2015. Seasonal and interannual variability of the marine bacterioplankton community throughout the water column over ten years. ISME J 9: 563–580.

32. Nelson CE. 2009. Phenology of high-elevation pelagic bacteria: the roles of meteorologic variability, catchment inputs and thermal stratification in structuring communities. ISME J 3: 13–30.

33. Kara EL, Hanson PC, Hu YH, Winslow L, McMahon KD. 2013. A decade of seasonal dynamics and co-occurrences within freshwater bacterioplankton communities from eutrophic Lake Mendota, WI, USA. ISME J 7: 680–4.

34. Yannarell AC, Kent AD, H. LG, Kratz TK, Triplett EW. 2003. Temporal Patterns in Bacterial Communities in Three Temperate Lakes of Different Trophic Status. Microb Ecol 46: 391–405.

35. Kent AD, Yannarell AC, Rusak JA, Triplett EW, McMahon KD. 2007. Synchrony in aquatic microbial community dynamics. ISME J 1: 38–47.

36. Kent AD, Jones SE, Lauster GH, Graham JM, Newton RJ, McMahon KD. 2006. Experimental manipulations of microbial food web interactions in a humic lake: Shifting biological drivers of bacterial community structure. Environ Microbiol 8: 1448–1459.

37. Rusak JA, Jones SE, Kent AD, Shade A, McMahon TD. 2009. Spatial synchrony in microbial community dynamics : testing among-year and lake patterns. Verh Internat Verein Limnol 30: 936–940.

38. Hug LA, Baker BJ, Anantharaman K, Brown CT, Probst AJ, Castelle CJ, Butterfield CN, Hernsdorf AW, Amano Y, Ise K, Suzuki Y, Dudek N, Relman DA, Finstad KM, Amundson R, Thomas BC, Banfield JF. 2016. A new view of the tree of life. Nat Microbiol 1: 1–6.

39. Gies EA, Konwar KM, Beatty JT, Hallam SJ. 2014. Illuminating microbial dark matter in meromictic Sakinaw Lake. Appl Environ Microbiol 80: 6807–6818.

40. Borrel G, Lehours AC, Bardot C, Bailly X, Fonty G. 2010. Members of candidate divisions OP11, OD1 and SR1 are widespread along the water column of the meromictic Lake Pavin (France). Arch Microbiol 192: 559–567.

41. Shade A, Kent AD, Jones SE, Newton RJ, Triplett EW, McMahon KD. 2007. Interannual dynamics and phenology of bacterial communities in a eutrophic lake. Limnol Oceanogr 52: 487–494.

42. Caporaso JG, Lauber CL, Walters WA, Berg-Lyons D, Huntley J, Fierer N, Owens SM, Betley J, Fraser L, Bauer M, Gormley N, Gilbert JA, Smith G, Knight R. 2012. Ultra-high-throughput microbial community analysis on the Illumina HiSeq and MiSeq platforms. ISME J 6: 1621–1624.

43. Amir A, McDonald D, Navas-Molina J, Kopylova E, Morton J, Xu Z, Kightley E, Thompson L, Hyde E, Gonzalez A, Knight R. 2017. Deblur Rapidly Resolves Single-Nucleotide Community Sequence Patterns. mSystems 2: 1–7.

44. Schloss PD, Westcott SL, Ryabin T, Hall JR, Hartmann M, Hollister EB, Lesniewski RA, Oakley BB, Parks DH, Robinson CJ, Sahl JW, Stres B, Thallinger GG, Van Horn DJ, Weber CF. 2009. Introducing mothur: Open-source, platform-independent, community-supported software for describing and comparing microbial communities. Appl Environ Microbiol 75: 7537–7541.

45. DeSantis TZ, Hugenholtz P, Larsen N, Rojas M, Brodie EL, Keller K, Huber T, Dalevi D, Hu P, Andersen GL. 2006. Greengenes, a chimera-checked 16S rRNA gene database and workbench compatible with ARB. Appl Environ Microbiol 72: 5069–5072.

46. Camacho C, Coulouris G, Avagyan V, Ma N, Papadopoulos J, Bealer K, Madden TL. 2009. BLAST plus : architecture and applications. BMC Bioinformatics 10: 1–9.

47. Wang Q, Garrity GM, Tiedje JM, Cole JR. 2007. Naive Bayesian Classifier for Rapid Assignment of rRNA Sequences into the New Bacterial Taxonomy. Appl Environ Microbiol 73: 5261–5267.

48. Lozupone C, Knight R. 2005. UniFrac : a New Phylogenetic Method for Comparing Microbial Communities UniFrac : a New Phylogenetic Method for Comparing Microbial Communities. Appl Environ Microbiol 71: 8228–8235.

49. De Cáceres M, Legendre P. 2009. Associations between species and groups of sites: indices and statistical inference. Ecology 90: 3566–3574.

